# Quantitative Systems Pharmacology Modeling Framework of Autophagy in Tuberculosis: Application to Adjunctive Metformin Host-Directed Therapy

**DOI:** 10.1101/2022.03.10.483882

**Authors:** Krina Mehta, Tingjie Guo, Robert Wallis, Piet H. van der Graaf, J.G. Coen van Hasselt

## Abstract

**Background:** Quantitative systems pharmacology (QSP) modeling of the host-immune response against Mtb can inform rational design of host-directed therapies (HDTs). We aimed to develop a QSP framework to evaluate the effects of metformin-associated autophagy-induction in combination with antibiotics.

**Methods:** A QSP framework for autophagy was developed by extending a model for host-immune response to include AMPK-mTOR-autophagy signalling. This model was combined with pharmacokinetic-pharmacodynamic models for metformin and antibiotics against Mtb. We compared the model predictions to mice infection experiments, and derived predictions for pathogen and host-associated dynamics in humans treated with metformin in combination with antibiotics.

**Results:** The model adequately captured the observed bacterial load dynamics in mice Mtb infection models treated with metformin. Simulations for adjunctive metformin therapy in newly diagnosed patients suggested a limited yet dose-dependent effect of metformin on reducing the intracellular bacterial load and selected pro-inflammatory cytokines. Our predictions suggest that metformin may provide beneficiary effects when overall bacterial load, or extracellular-to-intracellular bacterial ratio is low, either early after infection or late during antibiotic treatment.

**Conclusions:** We present the first QSP framework for HDTs against Mtb, linking cellular-level autophagy effects to disease progression. This framework may be extended to guide design of HDTs against Mtb.

## Introduction

The increasing burden of *Mycobacterium tuberculosis* (Mtb) infections is a major global health concern associated with approximately 1.5-2 million deaths annually [1]. Current first-line treatment to active tuberculosis (TB) disease, include a two-month intensive phase with rifampicin, isoniazid, pyrazinamide, and ethambutol (HRZE), followed by a four-month continuation phase with rifampicin and isoniazid (HR). A long treatment duration, common treatment failure, relapse, and emergence of multi-drug resistant Mtb strains are key challenges to successful TB treatment [2].

Host-directed therapies (HDT) aim to exploit the interplay between the pathogen and the host immune response [3,4]. HDTs are increasingly being studied for treatment against Mtb infections. One of the most studied HDT strategies to date is autophagy induction [5]. Autophagy is an intracellular catabolic process involving delivery of excessive or damaged cellular components, including bacteria, to lysosome for degradation to maintain homeostasis. The AMPK-mTOR signalling pathway is an important regulator of autophagy. Mtb activates phosphorylation of Akt, which stimulates phosphorylation of mTORC1. Activation of mTORC1 inhibits autophagy by phosphorylation of various autophagy-related proteins [6]. Preclinical studies have demonstrated involvement of mTOR signalling pathway in the host response to Mtb, suggesting its relevance as therapeutic target [6,7]. Therefore, metformin, an antihyperglycemic agent and mTORC1 inhibitor, has been proposed as potential HDT against Mtb [6,7]. Metformin adjunctive therapy in diabetic TB patients was found to be associated with an improved therapy success rate and lowered mortality [8,9]. The translation of experimental findings towards patients and subsequent optimal and rational design of HDT treatment strategies is challenged by the complex nature of the host-pathogen interactions [3].

Quantitative systems pharmacology (QSP) models aim to capture mechanistic details of the interactions between a biological system and pharmacokinetic-pharmacodynamic (PKPD) properties of a drug [10]. QSP-based characterization of drug-host-pathogen interactions may allow evaluation of expected treatment responses upon perturbation of specific targets, which may help to identify promising HDT targets and to evaluate different potential combination treatment strategies. Within the TB field, the mathematical modeling approaches have primarily focused on pharmacokinetic (PK) and pharmacodynamic (PD) modeling focusing mostly on design of antibiotics [11,12]. In addition, multiscale systems biology models of the host-immune response in response to Mtb infections have been developed[13,14]. The prior immune response model [13] have been combined with PKPD models of standard antibiotics to explore the impact of patient immune response on the treatment outcomes [15–17]. Any of these models did not include HDT relevant pathways; however, established a strong basis for further developing QSP framework to enable design of HDTs.

To guide design and development of HDTs, relevant HDT pathways must be added to the QSP framework. The autophagy-regulating AMPK/mTOR pathway represents an important factor for HDTs. There are currently no mathematical models available in literature describing mTOR signalling-mediated autophagy in TB. The objectives of this work were, (1) to develop a QSP framework of host-immune response including autophagy-mediated interactions, and (2) to evaluate the effects of metformin-associated autophagy-induction in combination with standard TB antibiotic treatment.

## Methods

The developed QSP framework (**Figure 1**) included, (1) PK models for standard antibiotics and metformin, (2) TB host-immune response model including PD effects of HRZE, and (3) autophagy model including PD effects of metformin. The QSP model development was facilitated by adaptations of various models presented in literature [17–19]. A set of ODEs describing dynamics of intra- and extra-cellular bacteria in host lungs as functions of time, and dynamics of immune-response components, such as macrophages, cytokines, and lymphocytes as functions of time and bacterial load, form the core of our QSP framework [18]. The core model was then linked to a model describing the dynamics of AMPK-mTOR signaling proteins leading to autophagy [19]. The interactions between Mtb and autophagy connects these two models. Moreover, the combined TB host-immune response-autophagy model was linked with models capturing concentrations-effect (PKPD) relationships of standard antibiotics and metformin.

**Figure 1.**
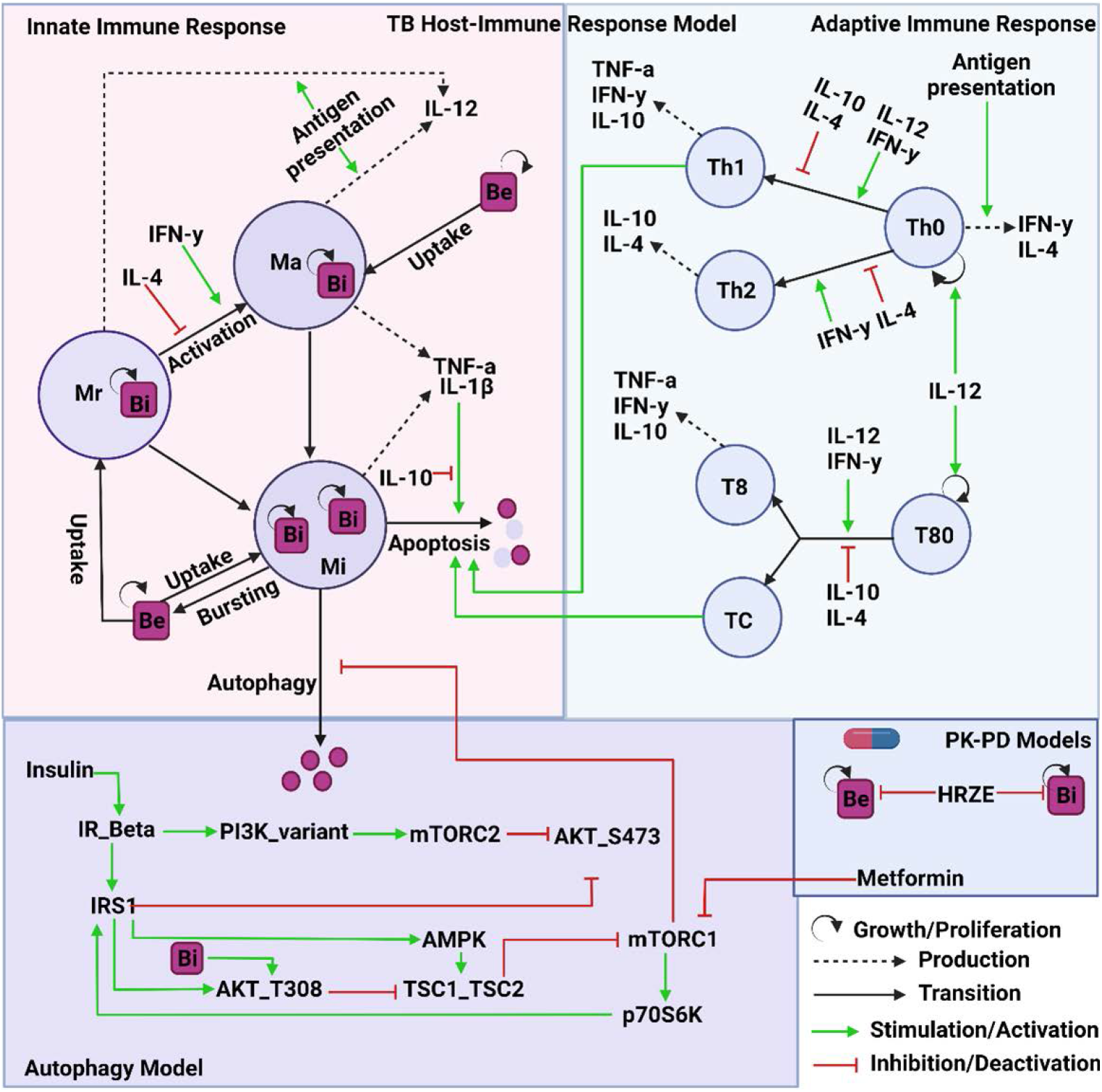
Combined TB-Autophagy QSP framework. The model captures the dynamics of host-immune response in the lungs because of Mtb infection. The model consists of various species of macrophages, lymphocytes, and the key cytokines involved in both innate and adaptive immune response against Mtb. The model includes the growth of Mtb as well as immune-mediated elimination of Mtb affecting the overall Mtb population. The immune-mediated bacterial killing include mainly cytokine- and lymphocytes-mediated apoptosis as well as autophagy. The model also consists of Mtb evasion mechanism, such as, induction of AMPK-mTOR pathway and inhibition of autophagy. Bi=intracellular Mtb, Be=extracellular Mtb, Ma=activated macrophage, Mi=infected macrophage, Mr=resident macrophage, T80=precursor-activated CD8+ T cells, T8=sub-class (IFN-y producing) of activated CD8+ T-cells, Tc=subclass (cytotoxic lymphocytes) of activated CD8+ T-cells, Th0=naïve T-cells, Th1=type 1 helper T-cells, Th2=type 2 helper T-cells.

### Model Development

The details of the model development process are provided in **S1**, and the key steps are presented below.

#### Pharmacokinetics

PK models of four antibiotics, HRZE, were reproduced from literature-based population pharmacokinetic (PopPK) models [20,21]. Plasma concentrations of HRZE following standard of care dosing were simulated using the PK models. HRZE intra- and extra-cellular lung concentrations were predicted by applying plasma to lung alveolar cells (AC) and plasma to lung epithelial lining fluid (ELF) ratios respectively obtained from literature [22]. To predict lung concentrations of metformin, we developed a minimal physiologically based pharmacokinetic model for metformin including a lung compartment [23] (**S1.1**).

#### TB Host-Immune Response and Pharmacodynamics of Standard Antibiotics

A published model that captured the dynamics of the host-pathogen interactions following Mtb infection was implemented [18]. This host-immune response model contained host-pathogen interactions in lungs, and included three population of macrophage (resting-, activated-, and infected-), various cytokines (IFN-γ, TNF-α, IL-10, IL-4, IL-12) and lymphocytes, as well as intra- and extra-cellular Mtb populations. An update was made to this model to add the turn-over of IL-1b and IL-1b-mediated bacterial elimination [24] (**Figure S1.2**).

We included two Mtb growth phases, fast and slow. The switch from fast to slow growth rates was empirically set to 21 days post-infection based on mice infection experimental results [25,26]. We used the slow phase bacterial growth rate estimates same as the growth rate values from the reproduced TB host-immune response model[18]. The growth rates for the initial fast phase were optimized using digitized data from mice Mtb infection experiment [25] (**Figure S1.2**). Bactericidal effects on intra- and extra-cellular bacterial population and bacteriostatic effects on growth rates of bacteria driven by intra- and extra-cellular lung concentrations of HRZE were reproduced from the literature [17].

#### Autophagy and Pharmacodynamics of Metformin

The AMPK-mTOR cell signalling network model from literature was reproduced [19]. This model captured the dynamics of key proteins involved in AMPK-mTOR signaling pathway and includes relative interactions between proteins involved in AMPK-mTOR signaling pathway, such as, insulin receptor substrate (IRS), class I phosphatidylinositol 3-kinases (PI3Ks), AMPK, mTORC1, and mTOR complex 2 (mTORC2). This model was updated to include various Mtb- and autophagy-related components. The updates can be categorized into: (1) the effect of Mtb Infection on autophagy inhibition due to activation of AMPK-mTOR signalling and (2) the effect of autophagy of Mtb elimination. Gene AKT3, a key upstream regulator of AMPK-mTOR signalling pathway, was found to be induced 1.38-fold in Mtb-infected vs. uninfected mice based on differential expression in lungs of Mtb infected vs. uninfected mice[6]. This ratio was added as a proportional scaling factor in the model on production of AKT to simulate the presence of Mtb and its impact on key down-stream proteins involved in AMPK-mTOR signalling, including mTORC1 (**S1.3**). Due to the limited data availability, time-course effects of progression of Mtb infection on autophagy is not included in the current model.

The effects of AMPK-mTOR signalling on autophagy was model using a direct effect saturable Emax model. Autophagy at time of Mtb infection was set to 100 % to represent healthy state prior to infection. Then, the percent inhibition of autophagy due to Mtb infection and subsequent AMPK-mTOR signalling activation modeled. Next, the autophagy model was combined with TB host-immune response model by introducing autophagy-mediated intracellular bacterial killing and autophagy-mediated extracellular to intracellular bacterial uptake. These processes were incorporated as first-order processes, and the parameters were informed by Mtb survival data from in vitro infection experiments with and without metformin treatment [27] (**S1.3**). The inhibitory effect of metformin on mTORC1 phosphorylation was incorporated using an indirect effect saturable Emax model, and the parameters were obtained from the literature [28,29].

### Model Evaluations

The combined TB-Autophagy QSP framework predictions were first compared to observed digitized lung bacterial load data from untreated and metformin-treated mice infected with Mtb [27]. To this end, the QSP model was scaled from humans to mice by applying lung volume differences between the species. To evaluate HRZE PKPD components of the combined QSP model, the predicted change in bacterial load over time after start of standard HRZE treatment was compared against reported values for TB patients [30–32].

### Sensitivity Analysis

High uncertainty existed in some parameters, especially for parameters related to autophagy model due to limited data availability. To further understand the impact of uncertainty in the parameters on model predictions, global uncertainty and sensitivity analysis using Latin hypercube sampling (LHS) and partial rank correlation coefficient (PRCC) method using 500 samples was performed [33,34] (**S3**). The outcome used in this analysis was predicted total bacterial load. All parameters, except the PK and PD parameters, were evaluated in the global uncertainty and sensitivity analysis. The parameters ranges used for LHS were the same as the previous model for TB host-immune response model components and were varied by 20% for autophagy-related components [18].

### Simulations of Metformin-Associated Autophagy Induction in Humans

Typical TB patient simulations were conducted using the QSP framework to predict the effects of autophagy induction with metformin on overall treatment outcome. Typical virtual TB patient simulations were performed using parameter values presented in **S2**. An initial extracellular Mtb inoculum of 100 bacteria was introduced at day 0 in all simulations. First, the simulations for TB disease progression, i.e., prior to the start of treatment, were conducted to evaluate consistency of the reproduced model with the original literature model. Next, simulations were performed to evaluate effects on bacterial load and on cytokine levels following standard TB therapy (2 months of HRZE + 4 months of HR) with and without adjunctive metformin treatment at three different dosing regimens starting at day 180 post-infection, i.e., upon diagnosis. Day 180 post-infection was selected as the approximate time to diagnosis and as such starting point for treatment based on prior model [17]. Metformin dosing regimen used in the simulations included 250 mg, 500 mg, and 1000 mg, all twice daily (BID) starting at day 180 post-infection. Additionally, to understand the effects of metformin on the TB disease progression in scenarios where diabetic patients would be receiving metformin for their glycemic control at the time of infection with TB [8,9], simulations were performed to predict the effects of 500 mg BID metformin treatment starting at day 1 post-infection.

### Software

All parameter optimization and model simulations were conducted in R and RStudio using nlmixr and RxODE packages [35]. Literature model for autophagy was converted from SBML file to ODEs in R using IQRsbml package [https://iqrsbml.intiquan.com/main.html].

## Results

The QSP framework included combined host-pathogen interactions model, AMPK-mTORC1 signalling pathway model including autophagy, and PK-PD models of HRZE and metformin (**Figure 1)**.

### The QSP Framework Simulations Recapitulate Observed In Vivo Response to Metformin

The model was evaluated by comparing predictions to the observed data. The model predictions for total bacterial load showed good agreement with observed digitized lung bacterial load data from untreated mice infected with Mtb, while the model showed slight underpredictions of the treatment effect in metformin-treated mice at day 35[6] (**Figure 2A)**. The simulations with standard TB therapy starting at day 180 post-infection in TB patients predicted previously reported change in bacterial load from baseline with standard TB therapy (2 HRZE + 4 HR) reasonably well (**Figure 2B**). Overall, these assessments suggested the reliability of the model for the objectives of this analysis.

**Figure 2.**
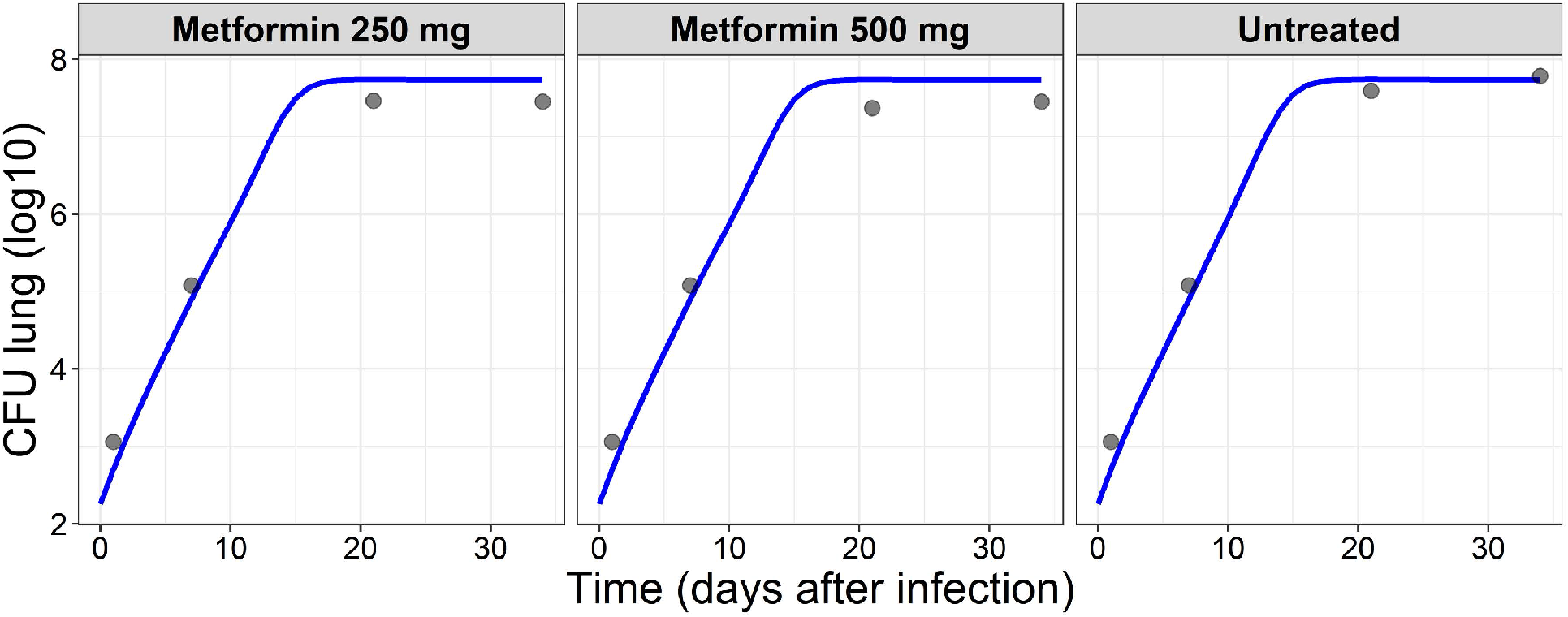
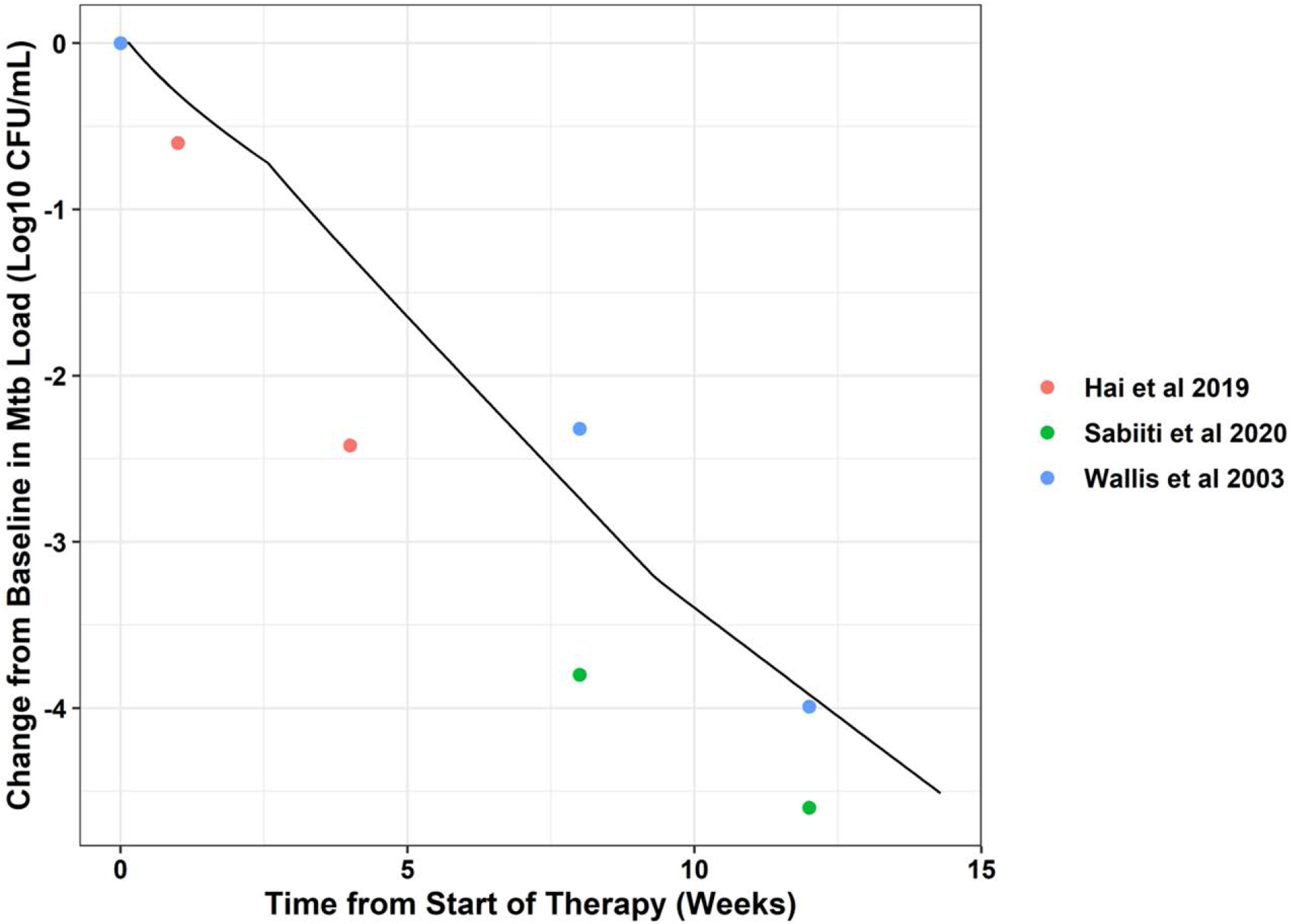
Time course of Predicted and Observed lung bacterial load: (A) Mtb-infected Mice Treated or Untreated with Metformin, (B) Tuberculosis Patients Treated with Standard Antibiotic Regimen. The model predictions for total bacterial load agree well with the observed mice data treated with or without metformin. Additionally, model predictions for effects of standard antibiotics therapy agree well with observed change in bacterial load from baseline data from TB patients. CFU=colony-forming unit, Metformin was administered daily from day 7 through 35 with six days on, one day off regimen, points represent observed and lines represent model predictions.

### Sensitivity Analysis Provides Insights into the Mechanistic Details of the Infection

The global uncertainty and sensitivity analysis suggested that the bacterial load was more sensitive to the parameters of host-pathogen interaction model compared to those of the autophagy model (**Figure S3**). In general, the host-pathogen interaction model parameters that correlated with the bacterial load the most were related to macrophage recruitment, macrophage activation or deactivation, phagocytosis, IFN-y production, or IL-1b- or FAS-FAS-mediated apoptosis. Most of these parameters were identified in the sensitivity analysis in the prior models too [18]. In the prior models, these parameters were obtained either from literature or were estimated using in vitro or mice experiments’ data and therefore, are considered relatively reliable. One parameter related to the autophagy model, AKT dephosphorylation rate, was found positively correlated with the bacterial load, and thus negatively correlated with infection control. This highlights the key role of Mtb evasion and inhibition of autophagy on disease progression.

This parameter was unchanged in the current model from the previous AMPK-mTOR signalling model. In the previous work, this parameter was estimated using experimental data from immunoblots and thus deem reliable. This sensitivity analysis given uncertainty in the parameters provide a thorough picture of the current state of the model.

### Metformin-Associated Autophagy Induction is Predicted to Provide Dose-Dependent Reduction in Intracellular Bacterial Load

The simulations for TB disease progression, i.e., prior to the start of treatment suggested that Mtb infection is predicted to reduce autophagy by 55% in a typical subject. Next, we compared the effects of standard TB antibiotic therapy (2 HRZE + 4 HR) with or without adjunctive metformin treatment on bacterial load and cytokine levels in a typical virtual TB patient. These simulations considered a typical scenario where the treatment was started upon diagnosis of TB, which was considered around day 180 after initial infection. Adjunctive metformin with 2 HRZE + 4 HR treatment was predicted to show apparent dose-dependent increase in autophagy-mediated intracellular bacterial elimination from the start of therapy (**Figure 3**). This increase in intracellular bacterial elimination with adjunctive metformin treatment however does not significantly affect extracellular and thus total bacterial elimination compared to standard antibiotics only treatment when total bacterial burden is high (>10^4^ mL^−1^). Some beneficial effect of adjunctive metformin treatment is predicted approximately 120 days after start of treatment when total bacillary load is relatively low. The simulations also showed metformin dose-dependent decrease in pro-inflammatory cytokines, IL-1b and IL-12, and dose-dependent increase in anti-inflammatory cytokine IL-10. Additionally, our results demonstrate there is not apparent difference in the levels of other cytokines, such as, IFN-y, TNF-α, and IL-4 in groups with or without adjunctive metformin treatment. Considering the total bacterial load and cytokine predictions we conclude that metformin treatment may provide some benefit in reducing intracellular bacterial load as well as in tissue damage caused by excessive pro-inflammatory cytokines in TB patients.

**Figure 3.**
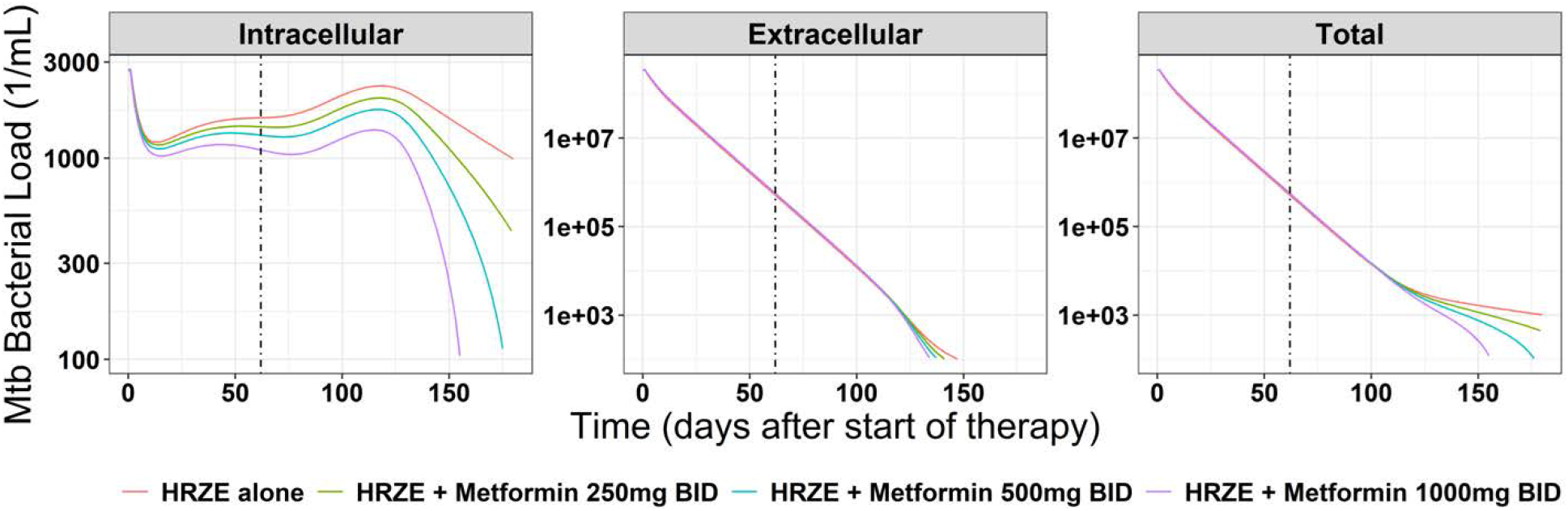
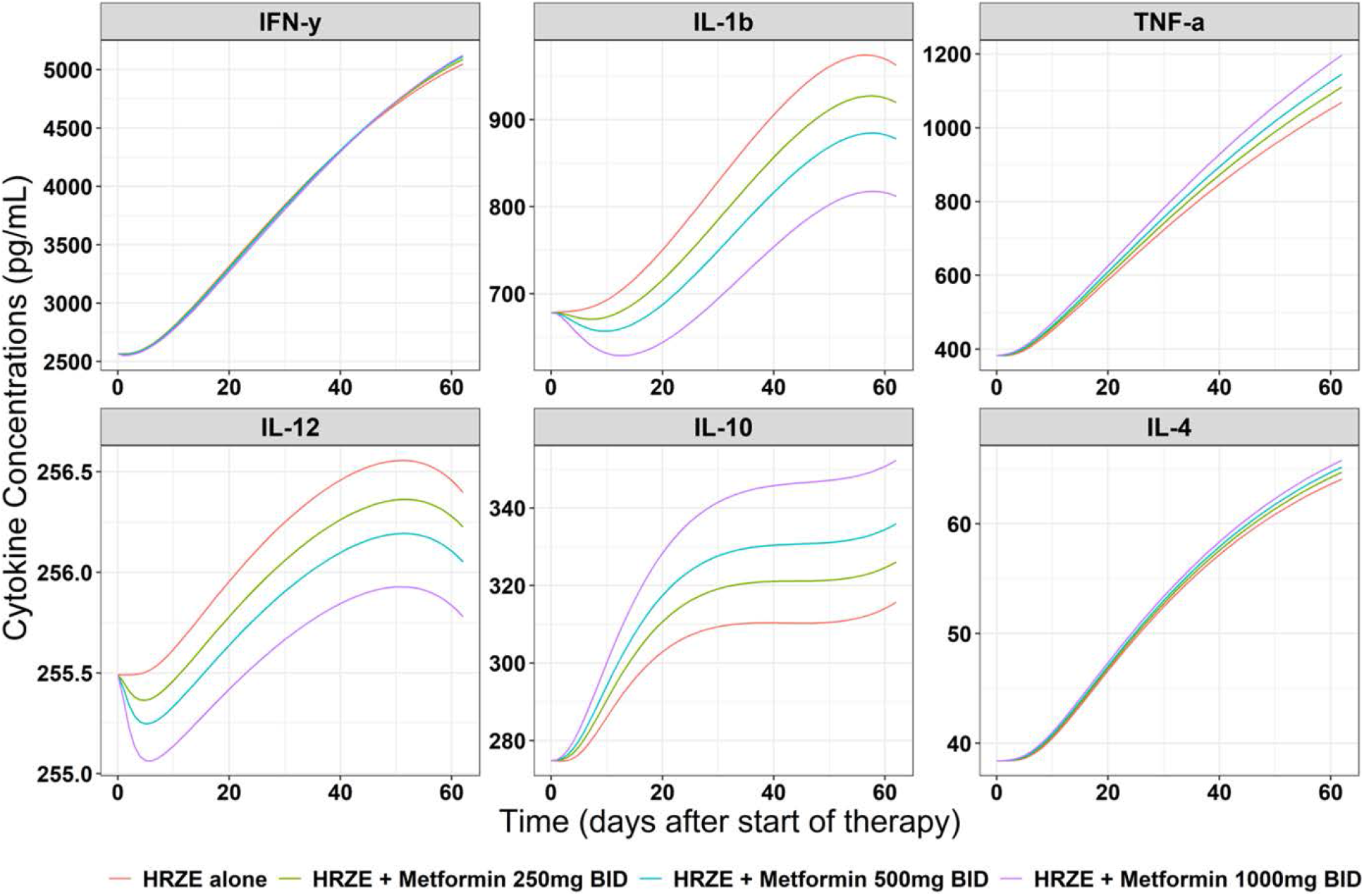
Typical Patient Simulations for Adjunctive Metformin to Standard Antibiotics Treatment at Various Dosing Regimen Starting at Day 180 Post-Infection: (A) Bacterial Load, (B) Cytokines. The simulations suggest dose-dependent effects of metformin on reduction of intracellular bacterial load and pro-inflammatory cytokines, IL-1b and IL-12. The reduction in intracellular bacterial elimination with adjunctive metformin treatment however does not significantly affect extracellular and thus total bacterial load compared to standard antibiotics only treatment when total bacterial burden is high. HRZE refers to 2 months of rifampicin, isoniazid, pyrazinamide, and ethambutol + 4 months of rifampicin and isoniazid regimen; vertical dashed line refers to end of 2 months regimen.

### Metformin May Delay Disease Progression in Diabetic TB Patientss

We assessed if metformin would delay TB disease progression if metformin was administered prior to TB diagnosis, i.e., in scenarios where diabetic patients would be receiving metformin for their glycemic control at the time of infection with Mtb (**Figure 4)**. In our simulations, metformin input was added at the same time as initial bacterial infection. We found that metformin use in diabetic TB patients would delay TB disease progression as assessed by intra-, extra-, and total-bacterial load. Lower levels of pro-inflammatory cytokines, IL-1b and IL-12, were also predicted in metformin-treated vs. no metformin-treated typical patient. Contrarily, slightly higher maximum levels of two pro-inflammatory cytokine, TNF-α and IFN-y, were predicted in metformin-treated vs. no metformin-treated typical patient. Overall, these simulations suggest some protective effects over tissue damage of metformin use in diabetic TB patients.

**Figure 4.**
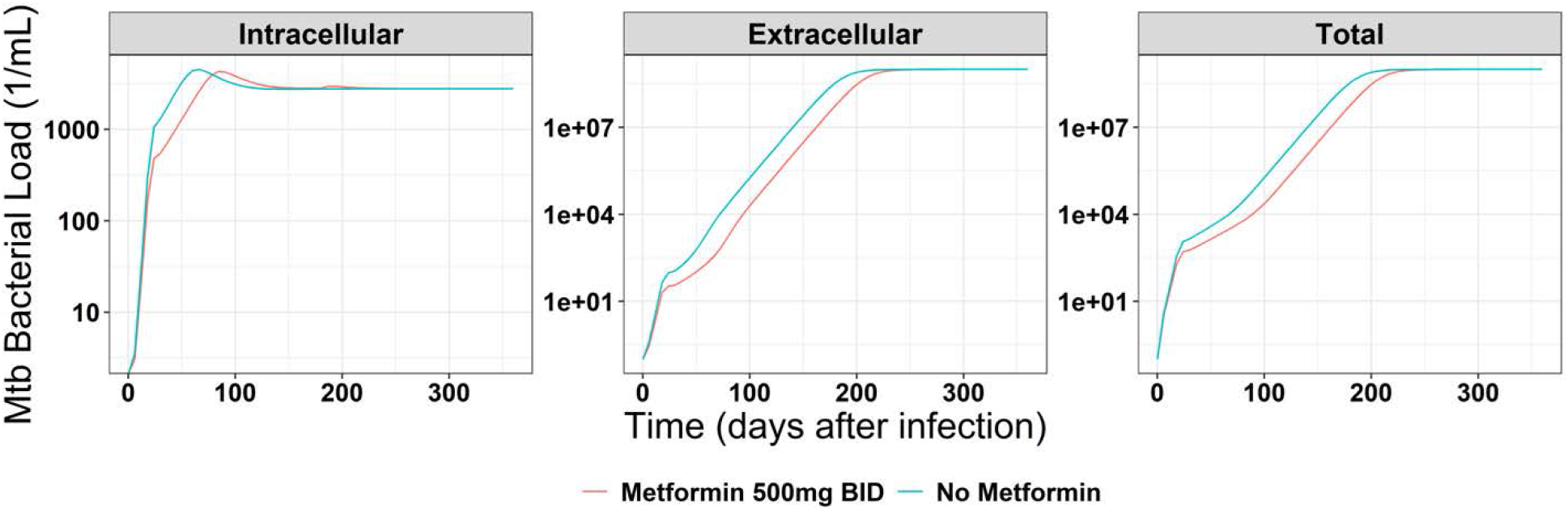
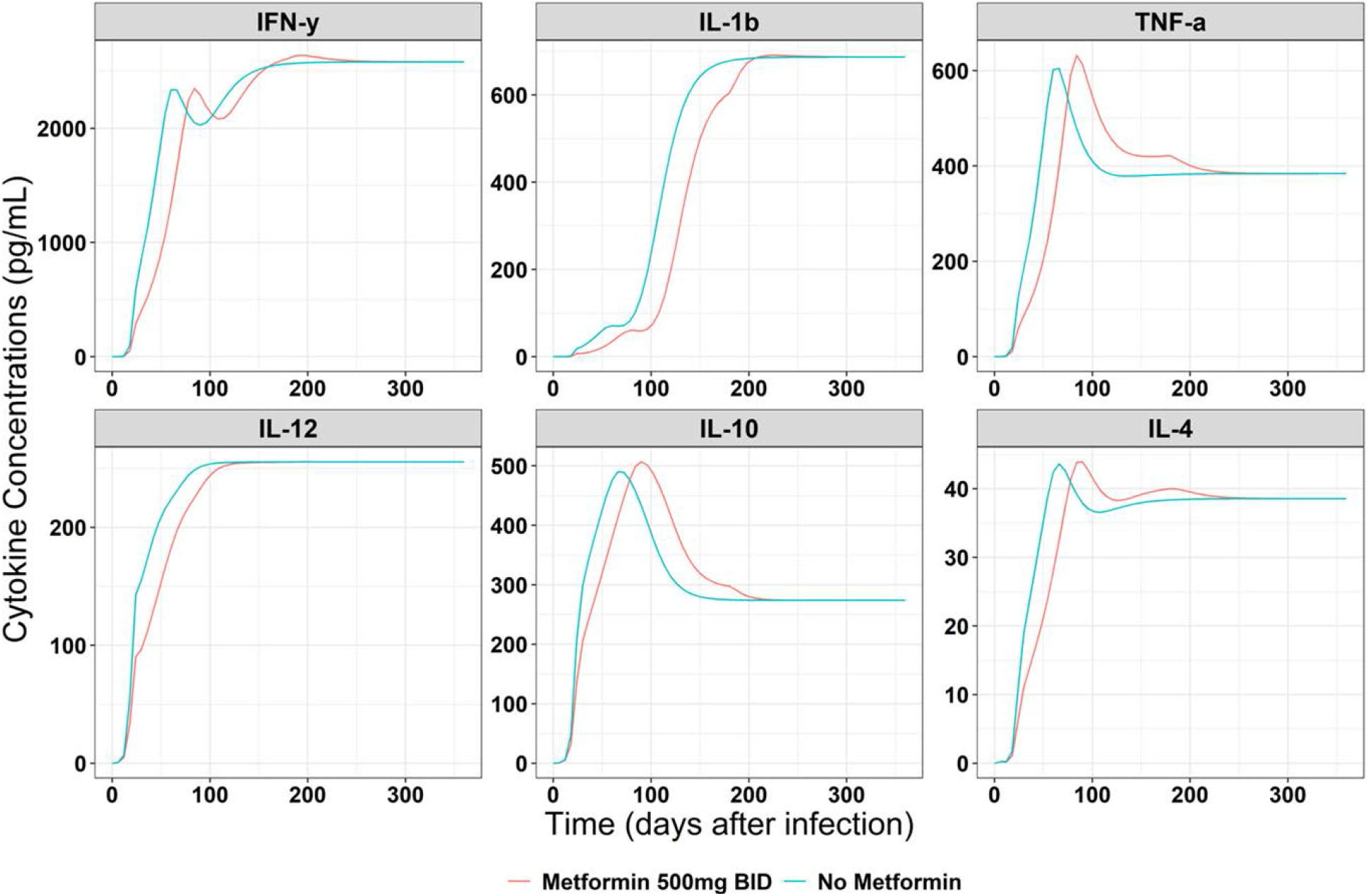
Typical Patient Simulations with or without 500 mg Twice Daily Metformin Starting at Day 1 Post-infection: (A) Bacterial Load, (B) Cytokines. The model predicted some benefits of metformin use in delaying the disease progression in virtual diabetic patients receiving metformin as compared to non-treated patients.

## Discussions and Conclusions

Here, we developed a first QSP framework for design and evaluation of HDTs focusing on autophagy. Our model was able to recapitulate results from an in vivo study evaluating metformin as a host-directed therapy in Mtb-infected mice (**Figure 2A**). We applied the framework to predict treatment effects of autophagy induction by metformin in a typical TB patient.

Our analysis identified a modest beneficial effect of adjunctive metformin treatment in a typical TB patient, approximately 120 days after start of treatment when total bacillary load is predicted to be relatively low (**Figure 3A**). The predictions suggested that overall effects of treatment with metformin would depend on extracellular-to-intracellular bacteria ratio, which may depend on the stage of infection. The model also predicted some benefits of metformin use in delaying the disease progression in virtual diabetic patients receiving metformin. Our results agree with the clinical reports where lowered mortality rates were reported in diabetic patients receiving metformin [8,9]. A key Mtb survival strategy depends on provoking a non-sterilizing immune response, allowing for Mtb to replicate beyond reach of most immune mechanisms. As part of host-pathogen interactions, granuloma formation limits Mtb growth, but also provide niche for replication by disseminating Mtb to other areas [36]. Metformin, and HDTs in general, may provide beneficiary effects early after initial infection, i.e., in newly infected TB household contacts, or late during treatment, i.e., after sputum has been sterilized but when small numbers of persisting bacteria are still present. In these scenarios, small changes in the survival of a rather small bacterial population may have a large effect on infection outcome, and future studies may consider evaluating this.

Our model provides relevant quantitative insight into the mechanistic details of factors contributing to autophagy-mediated bacterial elimination. Lack of predicted effects of metformin at doses up to 1000 mg BID on total bacterial load can also be attributed to its potency on AMPK-mTORC1-autophagy signaling and its distribution in lungs, in addition to extracellular-to-intracellular bacterial ratio. Previously, metformin dose-ranging study that evaluated effects of metformin at doses 100–10000 uM on Mtb survival in human monocyte-derived macrophages showed no increased Mtb survival at doses up to 500 uM. In the same study, approximately 4% reduction in total bacterial load on day 35 was noted in mice treated with 250 mg/kg and 500 mg/kg metformin daily from day 7-35 (6 days on, 1 day off)[6]. Our mPBPK model predicted mice lungs Cmax 668 uM and 1336 uM in 250 mg/kg and 500 mg/kg metformin dose groups, respectively. When these body weight-based doses of metformin that were evaluated in mice are compared to clinically feasible doses in humans (up to 1000 mg), predicted lungs Cmax in humans are approximately 10- to 15-fold lower than those predicted in mice. As such, it would be no surprise that our predictions showed no effects of metformin on the reduction of bacterial load. In fact, a recently completed clinical trial evaluating adjunctive metformin treatment to standard treatment in TB patients reported that metformin treatment did not reduce time to sputum conversion as compared to HRZE control arm [37].

The fine balance between levels of pro- and anti-inflammatory cytokines is an important attribute of the host-pathogen interactions affecting the overall disease outcome [3]. For example, high levels of pro-inflammatory cytokines may be associated with increased inflammation and tissue damage but can also reduce immune-mediated bactericidal activity [24,38]. A recent clinical trial reported that adjunctive metformin treatment to HRZE reduced inflammatory markers and reduced lung tissue damage [37]. Our model predicted reduced levels of pro-inflammatory cytokines, IL-1b and IL-12, and increased levels of anti-inflammatory cytokine, IL-10, while reducing intracellular bacterial load in a typical TB patient treated with adjunctive metformin therapy starting at time of TB diagnosis (**Figure 3B)**. Cumulatively, looking at cytokine predictions, it may be interpreted that adjunctive metformin treatment in TB patients may provide some benefit in reducing tissue damage caused by excessive pro-inflammatory cytokines.

The integrated QSP framework connects the complex intracellular process, autophagy, to disease outcome at organism-level. The model can be easily adapted to perform evaluations of other mTORC1 inhibitors mTORC1-independent autophagy inducers in the future using similar approach as ours. Some candidate drugs include, everolimus, statins, PI3K inhibitors, and tyrosine kinase inhibitor. The model can also facilitate in silico evaluations of perturbations of various proteins involved in autophagy and predict their effects on the outcome, as such enable target identification for optimal autophagy induction. For example, the model may be used in combination with screening assays to prioritize further development of potential HDTs.

Our model was built upon prior TB host-immune response model. We selected relatively simpler model given the objectives of this work [18]. In our model, we added the transition from fast- to slow- Mtb growth phase, which was set empirically based on mice infection experiments data. In humans, approximately 10% of Mtb infected individuals may develop active disease, of which primary infections account for 10% and reactivation of latent infection account for 90% [39]. Thus, majority of active TB cases follow a period of latency, which is not fully captured in the current model. More complex model is relevant for our primary objective, i.e., to evaluate different treatment scenarios in TB patients when bacterial load has already reached relatively high.

The model includes relative activity of the key proteins involved in the AMPK-mTOR-autophagy signalling, and however, do not consider total concentrations of these proteins. The original data-driven AMPK-mTOR model that was adapted in this work was developed using immunoblot data from HeLa cells, and therefore considered relative activity of the proteins [19]. Direct measurements of these are not available to date. However, the current approach of using relative activities of the AMPK-mTOR signalling proteins to evaluate their downstream effects on autophagy provide a useful alternative in absence of absolute proteins data.

To summarize, we developed a QSP framework for autophagy-inducing HDT by integrating a previously developed models for AMPK-mTOR signalling, host-pathogen interactions, and PKPD. We extended the framework to include autophagy to enable in silico evaluations of adjunctive metformin to antibiotics in TB patients. Our predictions suggest that metformin may provide some beneficiary effects when overall bacterial load, or extracellular-to-intracellular bacterial ratio is low, i.e., early after infection or late during antibiotics treatment. Overall, this is the first QSP that links cellular-level events affecting autophagy to disease progression model and may further be developed to guide HDT design and development for treatment of TB.

## Author Contributions

KM and JGC were involved in designing the analysis; KM performed the hands-on analysis and preparation of the first draft of the manuscript; TG performed quality checks of the model codes; All authors were involved in reviews and revisions of the manuscript.

## Funding

No funding was received for this work.

## Conflict of Interests

RSW is a co-investigator in a clinical study of metformin use in TB supported by NIH and a principal investigator of metformin use in TB supported by the EC Horizon 2020 program. All the other authors declare no conflict of interest.

## Acknowledgments

We thank Sud et al., Sonntag et al., and Fors et al., for making their model codes available in literature.

## Data Sharing Statement

All data used in the analysis were obtained from literature and source literatures are cited where applicable. The final model code used in the simulations is provided in the supplementary materials.

## Supplementary Material Titles

S1. Model Development Methods and the Final Model Code

S2. Parameter Values used in the Simulations

S3. Uncertainty and Sensitivity Analysis: Parameters Affecting Total Bacterial Load

S4. Typical TB Patient Simulations TB Disease Progression for Bacterial Load, Cytokines, and Key AMPK-mTOR-Autophagy Signalling Markers

